# Differences in systemic immune parameters in individuals with lung cancer according to race

**DOI:** 10.1101/2024.06.07.597754

**Authors:** Mitchell S. von Itzstein, Jialiang Liu, Hong Mu-Mosley, Farjana Fattah, Jason Y. Park, Jeffrey A. SoRelle, J. David Farrar, Mary E. Gwin, David Hsiehchen, Yvonne Gloria-McCutchen, Edward K. Wakeland, Suzanne Cole, Sheena Bhalla, Radhika Kainthla, Igor Puzanov, Benjamin Switzer, Gregory A. Daniels, Yousef Zakharia, Montaser Shaheen, Jianjun Zhang, Yang Xie, David E. Gerber

## Abstract

**Introduction:** Racial and ethnic disparities in the presentation and outcomes of lung cancer are widely known. To evaluate potential factors contributing to these observations, we measured systemic immune parameters in Black and White patients with lung cancer.

**Methods:** Patients scheduled to receive cancer immunotherapy were enrolled in a multi-institutional prospective biospecimen collection registry. Clinical and demographic information were obtained from electronic medical records. Pre-treatment peripheral blood samples were collected and analyzed for cytokines using a multiplex panel and for immune cell populations using mass cytometry. Differences between Black and White patients were determined and corrected for multiple comparisons.

**Results:** A total of 187 patients with non-small cell lung cancer (Black, 19; White, 168) were included in the analysis. There were no significant differences in baseline characteristics between Black and White patients. Compared to White patients, Black patients had significantly lower levels of CCL23 and CCL27, and significantly higher levels of CCL8, CXCL1, CCL26, CCL25, CCL1, IL-1 b, CXCL16, and IFN-γ (all *P*<0.05, FDR<0.1). Black patients also exhibited greater populations of non-classical CD16+ monocytes, NKT-like cells, CD4+ cells, CD38+ monocytes, and CD57+ gamma delta T cells (all *P*<0.05).

**Conclusions:** Black and White patients with lung cancer exhibit several differences in immune parameters, with Black patients exhibiting greater levels of numerous pro-inflammatory cytokines and cell populations. The etiology and clinical significance of these differences warrant further evaluation.

---

The incidence, presentation, and outcomes of many cancers differ according to patient race. For instance, Black patients have greater incidence and mortality from prostate cancer.(1) Compared to White patients, Black individuals develop lung cancer after less smoking, at younger age, at more advanced stage, and have decreased survival.(2)

To evaluate these observations further in the immuno-oncology era, studies have also evaluated immune parameters according to patient race and ethnicity. IL-6 and CRP levels are elevated in healthy Black populations compared to White populations.(3) Differences also exist within oncology populations. For instance, Black patients with prostate cancer have higher levels of CXCL2 and CXCL5 than do White patients.(4) In resected lung cancer specimens, cell proliferation pathways and macrophage subtypes differed between Black and White patients.(5)

Given the persistent racial health disparities in lung cancer presentation and outcomes and the growing awareness of differences in inflammatory markers across populations, we analyzed systemic cytokines and immune cell populations in White and Black patients with NSCLC.

## Methods

### Study protocol and clinical data

This was a prospective multi-institutional registry study approved by the University of Texas Southwestern Institutional Review Board (IRB #STU 082015–053) and the IRBs of all participating centers. Patients provided written informed consent prior to undergoing any study-specific procedures.

Eligible patients had a diagnosis of cancer and were planned for but had not yet initiated ICI-based therapy, including concurrent and sequential combination regimens. Enrolled individuals underwent collection of clinical data and collection of pretreatment peripheral blood samples.

For this analysis, we analyzed patients with NSCLC of any stage who self-identified as non-Hispanic Black or non-Hispanic White.

### Biospecimen processing and cytokine analysis

As previously described,(6) we used the Bio-Plex Pro Human Chemokine 40-plex Panel (Bio-Rad Laboratories, Hercules, California) on a Luminex 200 System to measure plasma cytokines. **Supplemental Table 1** lists the cytokines included in the panel. Cytokine concentrations were transformed on a log_2_ scale and batch-corrected using the ComBat parametric empirical Bayes framework.(7)

### Cytometry by time of flight (CyTOF)

Cryopreserved peripheral blood mononuclear cells (PBMCs) were thawed and stained with a panel of 36 antibodies (metal isotope-labeled conjugates, Maxpar Direct Immune Profiling Assay Panel by Standard BioTools). We analyzed cells on a Helios mass cytometer (Standard BioTools). Data were normalized and analyzed with gating on CD45+ cells using the cloud-based computational platform OMIQ.ai (Dotmatics Software Company). We identified cluster immune phenotypes following standard immunophenotyping for the Human Immunology Project.

### Statistical analysis

We used chi-squared tests, Fisher’s exact tests, and Mann-Whitney U to assess for associations between case characteristics and race. Log_2_transformed and batch-corrected cytokine values were compared between the two groups using the Mann-Whitney U test. To account for multiple comparisons, the Benjamini–Hochberg procedure was applied to evaluate false discovery rates. All analyses were conducted with R (v4.1.3) or GraphPad Prism 10.2.3. We defined significance as *P*<0.05 and for cytokine analysis *P*<0.05 and FDR<0.1.

### Pathway analysis

We used the online tool string-db.org to analyze pathway interactions among cytokines exhibiting significant differences between Black and White patients. Both Gene Ontology (biological process) and Pathways (Kyoto Encyclopedia of Genes and Genomes (KEGG) Pathways and wikiPathways) were analyzed.

## Results

A total of 187 patients with non-small cell lung cancer (Black, 19; White, 168) were identified for this study. Among these individuals, 57 % were male, 65% were greater than 65 years old, and 81% had stage IV disease. Additional case characteristics are shown in **Table 1**. There were no significant differences between White and Black patient characteristics, although there was a near-significant trend towards earlier stage disease in Black patients (*P*=0.06).

**Table 1.** Clinical characteristics in the overall cohort and according to race.

| **Characteristic**<br>**n (%)** | **Overall**<br>**(N=187)** | **White**<br>**(N=168)** | **Black**<br>**(N=19)** | ***P* value** |
| --- | --- | --- | --- | --- |
| Age, years |  |  |  | 0.54 |
| ≥65 | 121 (65) | 107 (64) | 14 (74) |  |
| <65 | 66 (35) | 61 (36) | 5 (26) |  |
| Sex |  |  |  | 0.76 |
| Male | 107 (57) | 95 (57) | 12 (63) |  |
| Female | 80 (43) | 73 (43) | 7 (37) |  |
| Cancer stage |  |  |  | 0.06 |
| IV | 152 (81) | 140 (83) | 12 (63) |  |
| <IV | 35 (19) | 28 (17) | 7 (37) |  |
| Histology |  |  |  | 1 |
| Squamous | 39 (21) | 35 (21) | 4 (21) |  |
| Non-squamous | 148 (79) | 133 (79) | 15 (79) |  |
| Smoking status |  |  |  | 0.62 |
| Former | 118 (73) | 102 (71) | 16 (84) |  |
| Current | 19 (12) | 18 (13) | 1 (5) |  |
| Never | 25 (15) | 23 (16) | 2 (11) |  |
| Pack years |  |  |  | 0.53 |
| Median (IQR) | 30 (15-50) | 34 (14-50) | 25 (19-44) |  |
| BMI |  |  |  | 0.32 |
| <18.4 | 5 (3) | 5 (3) | 0 (0) |  |
| 18.5-24.9 | 62 (39) | 55 (38) | 7 (39) |  |
| 25-29.9 | 57 (35) | 53 (37) | 4 (22) |  |
| >30 | 37 (23) | 30 (21) | 7 (39) |  |

\*Categories that do not contain 187 cases represent missing data.

Cytokine levels were available for all 187 patients. Among the 40 cytokines analyzed, 35 (88%) had results that met technical criteria for inclusion in the study. Of these cytokines, 10 (29 %) exhibited statistically significant differences between Black and White patients (**Figure 1**). For eight of the cytokines with significant differences between races (80%), levels were higher in black patients, with CCL23 and CCL27 representing the only exceptions. **Supplemental Table 2** displays levels of all 35 included cytokines according to race.

**Figure 1.**
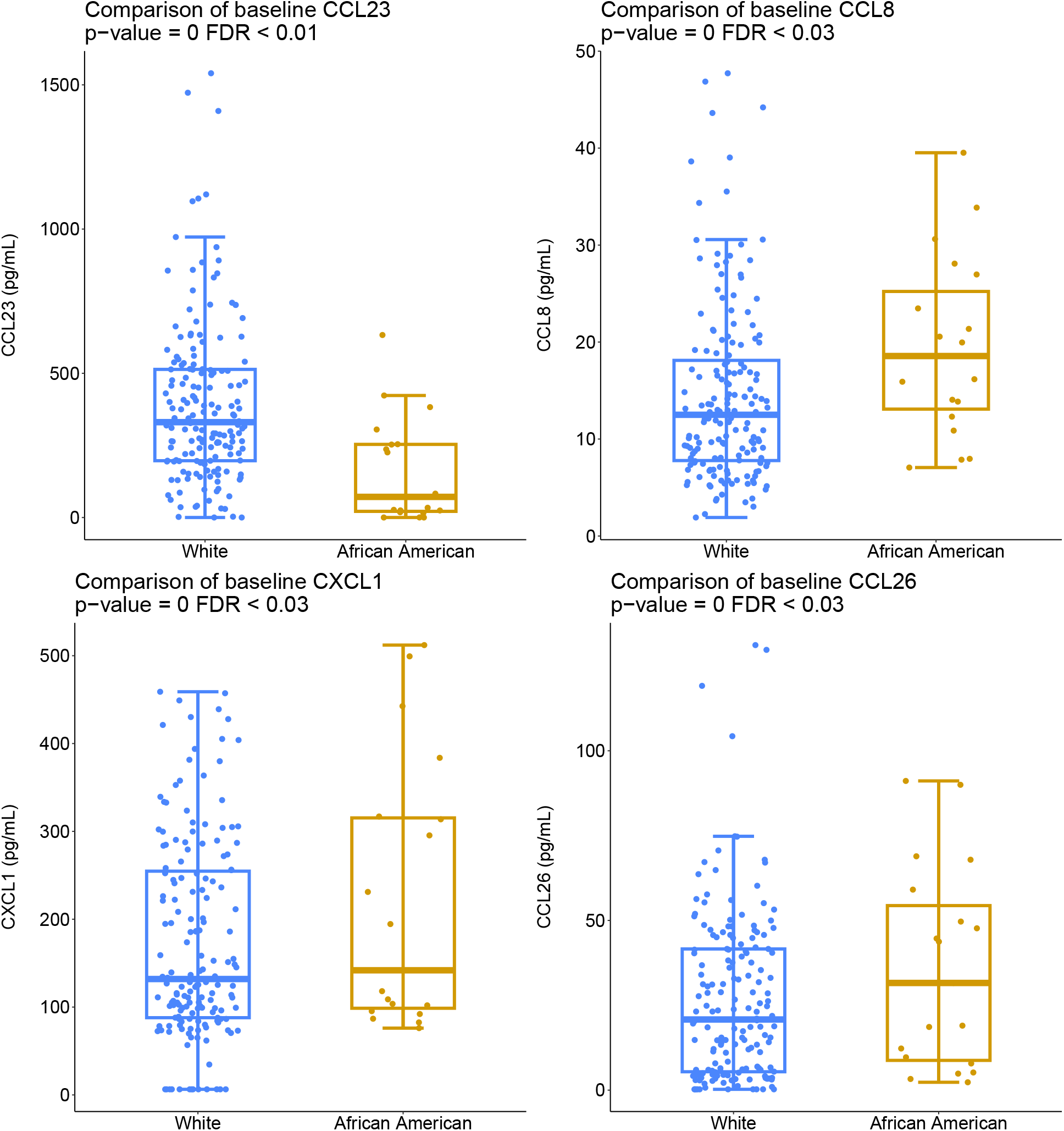
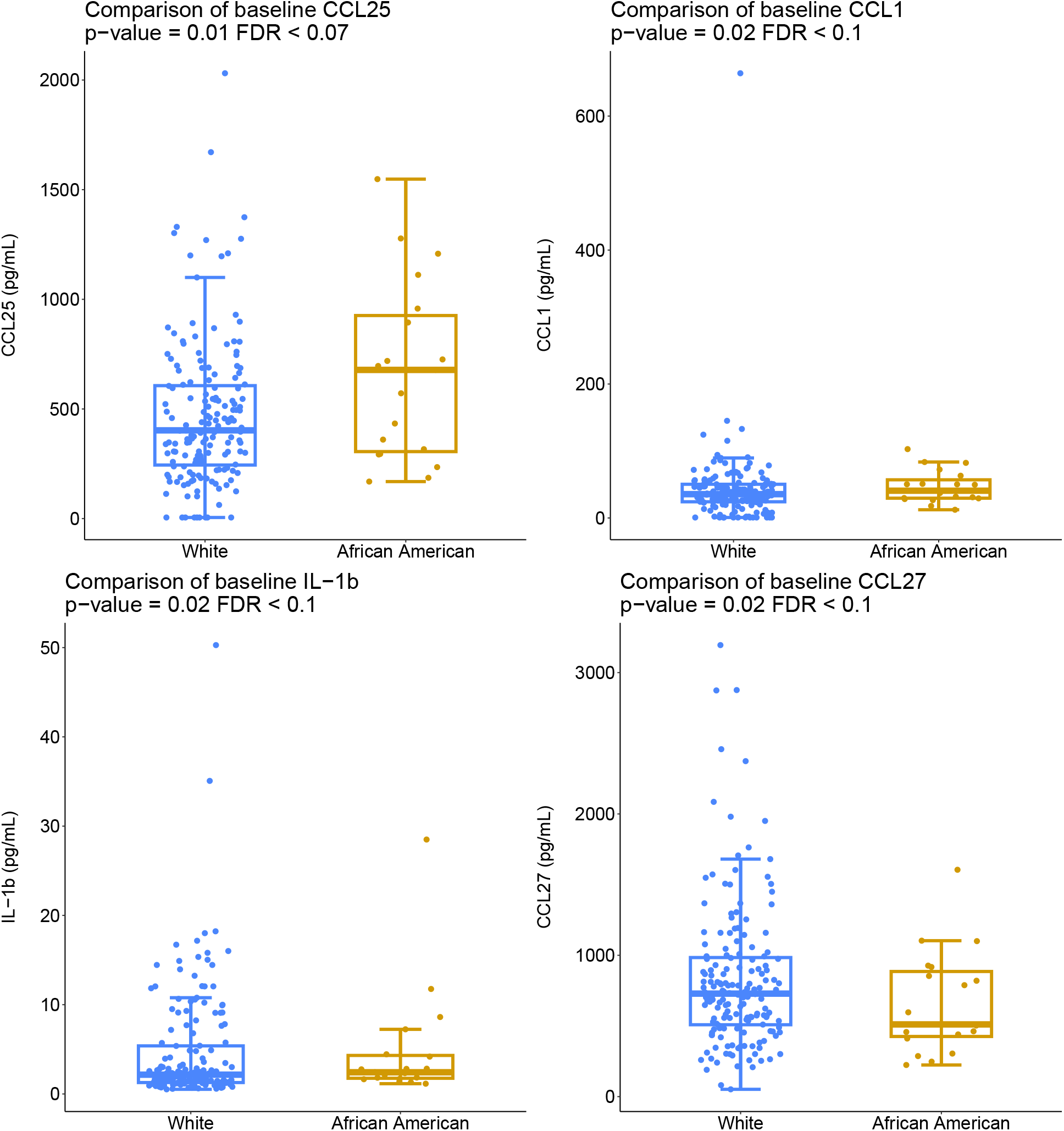
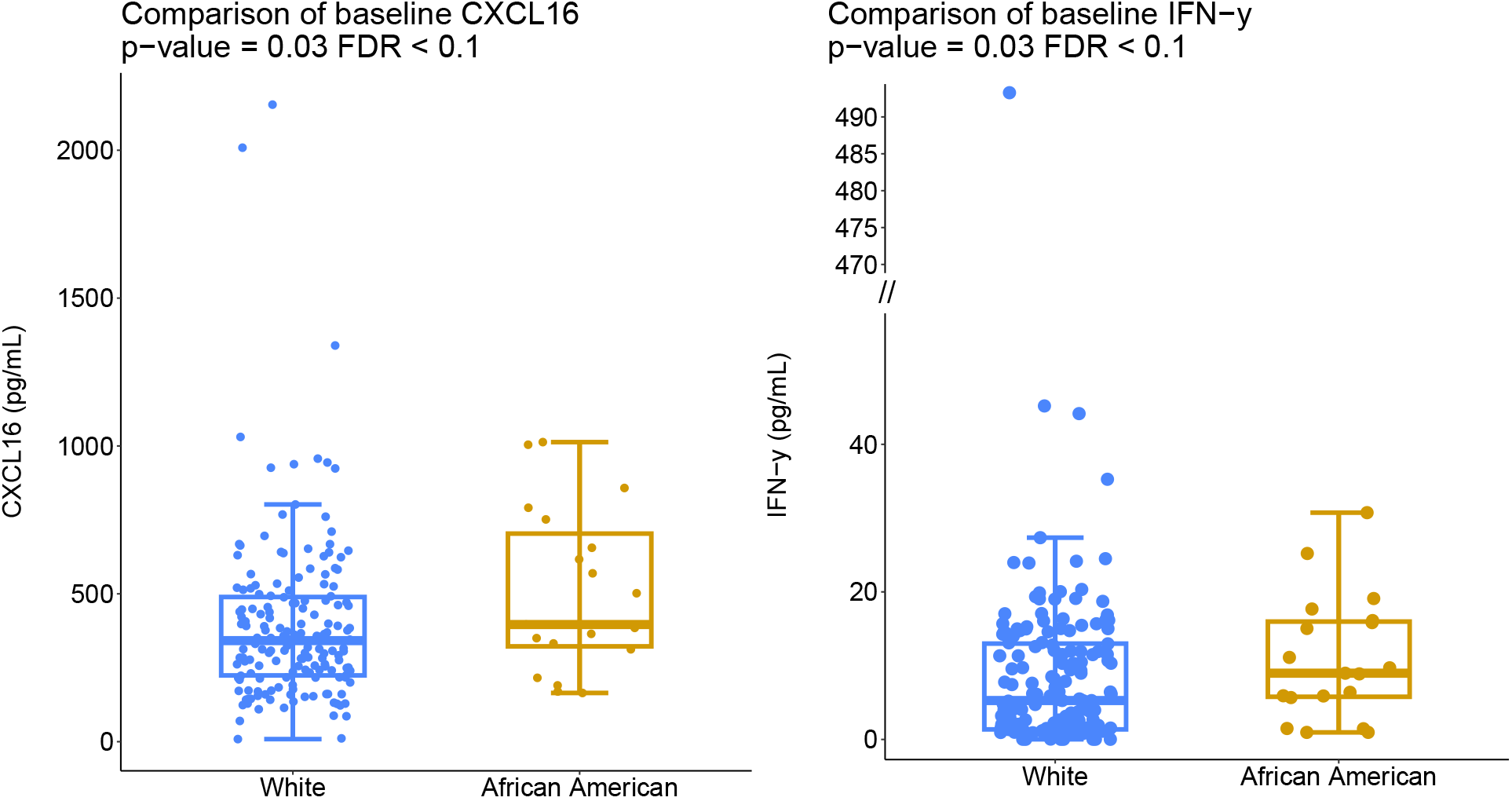
Cytokine analysis. Ten of 35 analyzed cytokines exhibited significant differences (*P*<0.05 and FDR<0.1) between Black and White populations.

Pathway analysis of the 10 cytokines with significant differences (P<0.05; FDR<0.1) indicated strong associations with five inflammatory signaling pathways, including IL-17, NF-kappa B, TNF, and NOD-like receptor signaling pathway (**Supplemental Figure 1**).

We analyzed PBMCs by CyTOF for 48 (Black, 16; White, 32) patients (26 %). Among 62 identified immune cell clusters, seven (11 %) had statistically significant differences between races (**Figure 2** and **Supplemental Figure 2**), all of which were elevated in black patients (*P* <0.05) (**Figure 2**). These included monocytes, natural killer (NK) T cell like (NKT-like) cells, CD4+, and gamma-delta T cells (γδT).

**Figure 2.**
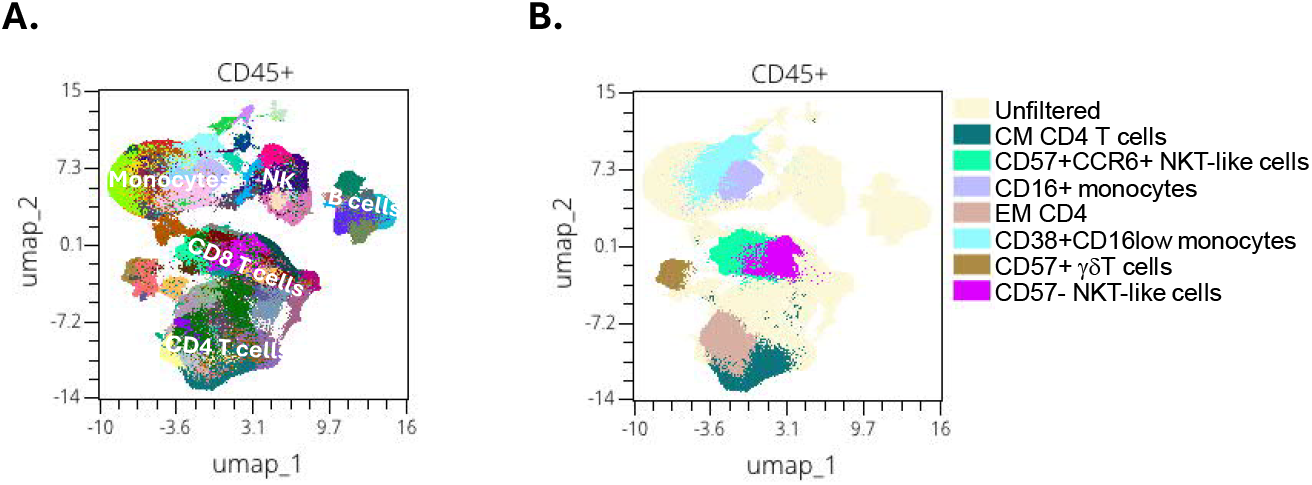
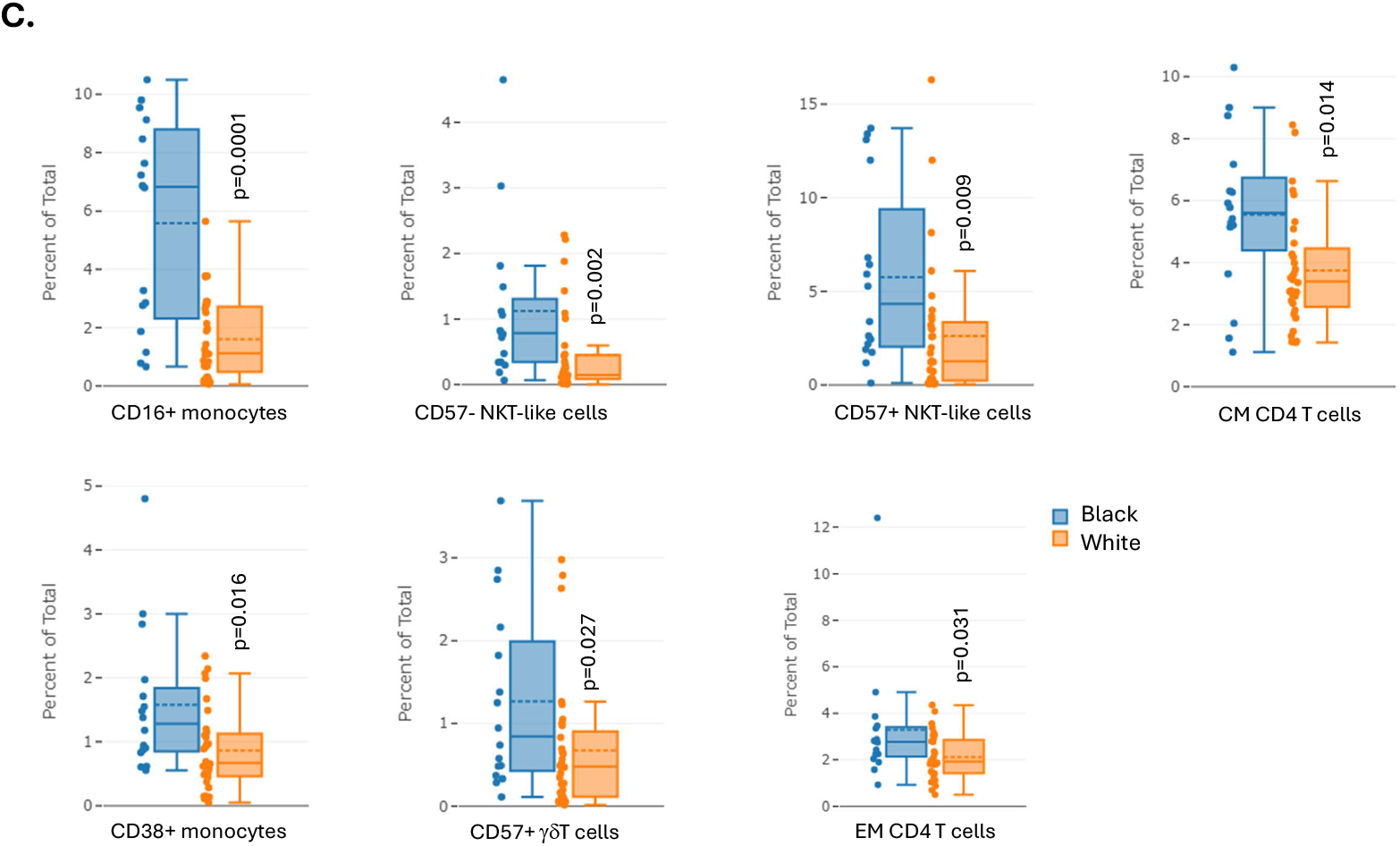
Immune cell profiling by cytometry by time of flight (CyTOF) analysis. ***(A)*** Uniform manifold approximation and projection (UMAP) showing 62 clusters (with major cell populations shown) identified by PhenoGraph-guided metaclustering of CD45+ cells in peripheral blood mononuclear cells (PBMCs). An equal number of cells per sample (38,614 cells / sample) in a total of 48 Black and White patients to create the UMAP map. ***(B)*** UMAP demonstrating 7 of 62 clusters (11%) with significant differences between Black and White populations. CM, central memory; EM, effector memory; NKT, natural killer T. ***(C)*** Immune cell subsets with significant differences between races. Solid lines designate median; dotted lines designate mean. Bars represent maximum and minimum values. Dots adjacent to boxes represent individual samples. *P* values by Mann-Whitney test. CM, central memory; EM, effector memory; NKT, natural killer T.

## Discussion

Immune responses play central roles in cancer development and progression, as well as chronic stressors associated with racial and ethnic disparities. We observed significant differences between Black and White patients for about 30 % of examined cytokines/chemokines and about 10 % of examined immune cell populations.

For almost all parameters, pro-inflammatory cytokine levels were higher in Black persons. Notably, almost all observed racial differences also appeared to have favorable prognostic implications for White individuals.The cytokine with substantially higher levels among White patients was CCL23, which in cancer tissue, correlates with the presence of macrophages, as well as PD-1, CTLA-4, TIGIT, LAG-3, and TIM-3 immune checkpoints,(8) and is associated favorable outcomes in immunotherapy-treated patients with lung cancer.(9) Conversely, CCL8, CXCL1, and CCL26—all significantly higher in Black patients in the present study—promote tumorigenesis and cancer progression in various malignancies.(10-12)

Results from the analysis of immune cell populations appear to correlate with our cytokine findings. We observed higher levels of non-classical monocytes and NKT-like cells in Black patients. Non-classical (CD16+) monocytes have potent inflammatory properties, serving as the primary producers of IL-1 β, which was elevated in Black compared to White patients. Among other cytokines, activated NKT cells produce IFN-γ,(13) which was elevated in Black patients.

The effect of these cells and their cytokines is to create a pro-inflammatory state. Further examination of these cellular findings is warranted as higher macrophage or CD4+ T cells can lead to a tolerant tumor microenvironment that does not allow immune mediated rejection. In contrast, NK cells and gamma-delta T cells are associated with cancer cell rejection and killing. Therefore, future efforts to examine subsets of these cell types to determine their transcriptional phenotype to infer activity are warranted.

Our observations correspond to findings in other populations. For instance, Black patients with prostate cancer have lower levels of CCL23 than do White patients with prostate cancer.(4) However, in a population study examining inflammatory markers associated with *risk* of lung cancer in Black and White individual, IFN-γ was associated with lung cancer risk in both races.(14) Among the five markers uniquely elevated in black patients, the three included in our dataset (IL-10, MCP-4/CCL13, and MIP-1/CCL3) were not associated with race in the present study. While our study focused on a different research question (comparing immune parameters in patients with existing lung cancer according to race), reasons for these differences are unclear. Our study expands on this prior study by incorporating a more extensive cytokine multiplex panel, additional analysis of circulating cell populations, and computational biological pathway analyses. Importantly, several markers identified in our and other studies in oncology populations have also been linked to other exposures. As an example, experiencing racial discrimination is associated with increased levels of pro-inflammatory cytokines (IL-1b, IL-6, IL-8, IL-10, TNF-α, and IFN-γ) in Black adolescents.(15)

This study only focused on baseline inflammatory and immune cell characteristics, but future studies could examine how these change with treatment. Especially in the context of ICI therapy, immune parameters may predict favorable responses or potential for immune-related adverse events. Comparing absolute differences in inflammatory/ immune cell levels and changes from baseline to follow up (especially longitudinally in the same patients) will provide deeper insight into the dynamic nature of the host immune system during ICI therapy for lung cancer.

A key limitation of this study is the limited number of Black patients, which in part reflects our inability to enroll patients at clinical sites providing care to large numbers of under-represented minorities during the COVID pandemic. We also lack information on socioeconomic status, which may influence inflammatory states and immune markers. While all blood samples were collected prior to ICI initiation, depending on the treatment regimen, other therapies such as radiation therapy may have been administered prior to blood collection. Due to the relatively small size of the study, we were not able to perform subgroup analyses according to other clinical characteristics. Strengths of the study include multi-center participation and inclusion of immune cell populations and pathway analysis in addition to cytokines.

Despite a relatively small sample size, this study demonstrates a more pro-inflammatory cytokine or cellular phenotype for Black patients with lung cancer. Although Black patients tended to have earlier-stage cancer in our cohort, almost all the immune parameter differences identified in Black patients are generally associated with worse outcomes. Further studies investigating the etiology and modulation of these observations are warranted.

---

**Supplemental Table 1.** Cytokines included in the analysis.

| **Total** | **Passed Quality Control** |
| --- | --- |
| 6Ckine/ CCL21 | 6Ckine/ CCL21 |
| BCA-1/ CXCL13 | BCA-1/ CXCL13 |
| CTACK/ CCL27 | CTACK/ CCL27 |
| ENA-78/ CXCL5 |  |
| Eotaxin/ CCL11 | Eotaxin/ CCL11 |
| Eotaxin-2/ CCL24 | Eotaxin-2/ CCL24 |
| Eotaxin-3/ CCL26 | Eotaxin-3/ CCL26 |
| Fractalkine/ CX3CL1 | Fractalkine/ CX3CL1 |
| GCP-2/ CXCL6 | GCP-2/ CXCL6 |
| GM-CSF |  |
| Gro-a/ CXCL1 | Gro-a/ CXCL1 |
| Gro-b/ CXCL2 | Gro-b/ CXCL2 |
| I-309/ CCL1 | I-309/ CCL1 |
| IFN-γ | IFN-γ |
| IL-1b | IL-1b |
| IL-2 | IL-2 |
| IL-4 | IL-4 |
| IL-6 |  |
| IL-8/ CXCL8 | IL-8/ CXCL8 |
| IL-10 | IL-10 |
| IL-16 | IL-16 |
| IP-10/ CXCL10 | IP-10/ CXCL10 |
| I-TAC/ CXCL11 | I-TAC/ CXCL11 |
| MCP-1/ CCL2 | MCP-1/ CCL2 |
| MCP-2/ CCL8 | MCP-2/ CCL8 |
| MCP-3/ CCL7 |  |
| MCP-4/ CCL13 | MCP-4/ CCL13 |
| MDC/ CCL22 | MDC/ CCL22 |
| MIF | MIF |
| MIG/ CXCL9 | MIG/ CXCL9 |
| MIP-1a/ CCL3 | MIP-1a/ CCL3 |
| MIP-1d/ CCL15 | MIP-1d/ CCL15 |
| MIP-3a/ CCL20 | MIP-3a/ CCL20 |
| MIP-3b/ CCL19 | MIP-3b/ CCL19 |
| MPIF-1/ CCL23 | MPIF-1/ CCL23 |
| SCYB16/ CXCL16 | SCYB16/ CXCL16 |
| SDF-1a+b/ CXCL12 | SDF-1a+b/ CXCL12 |
| TARC/ CCL17 |  |
| TECK/ CCL25 | TECK/ CCL25 |
| TNF-α | TNF-α |

**Supplemental Table 2.**
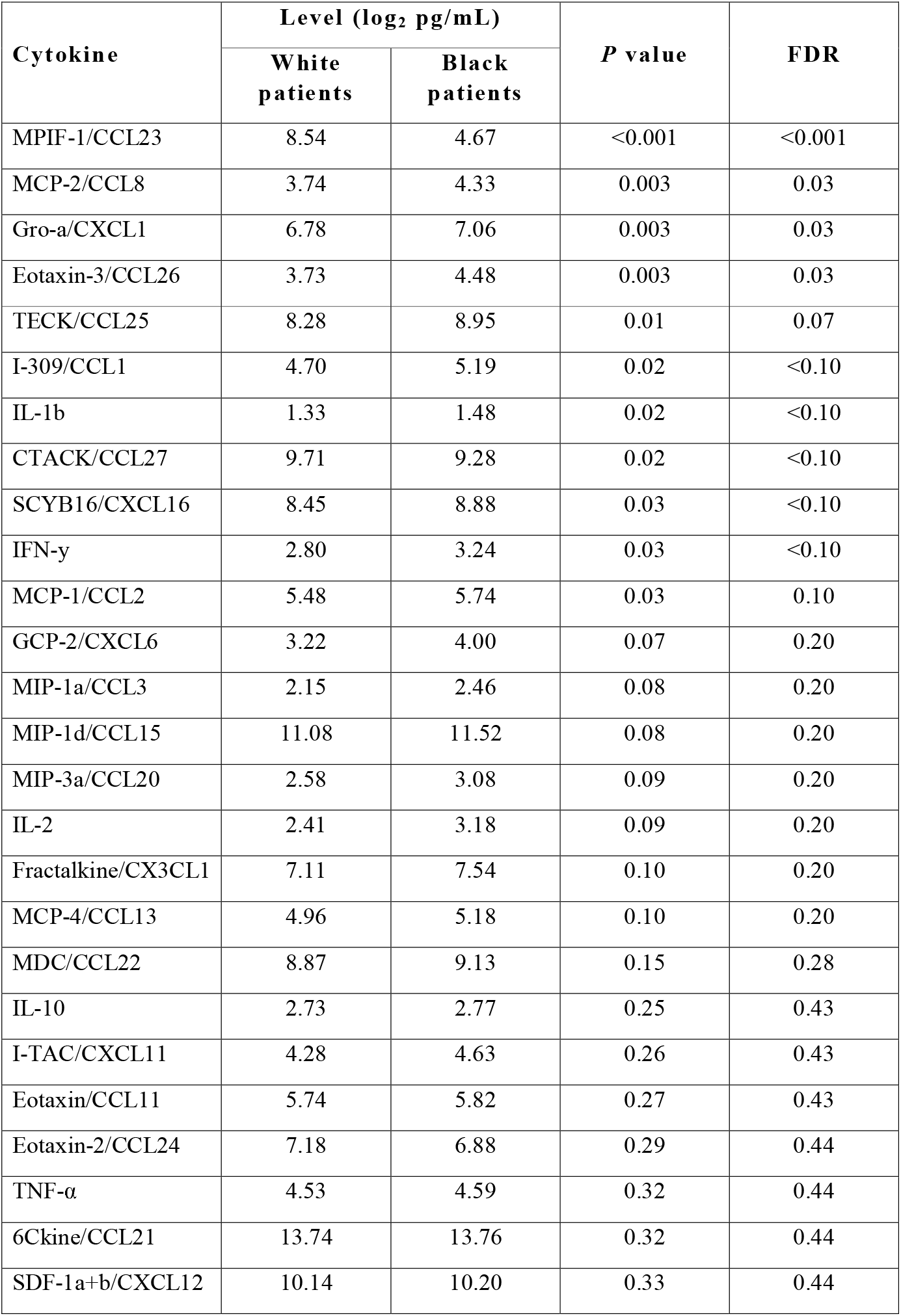
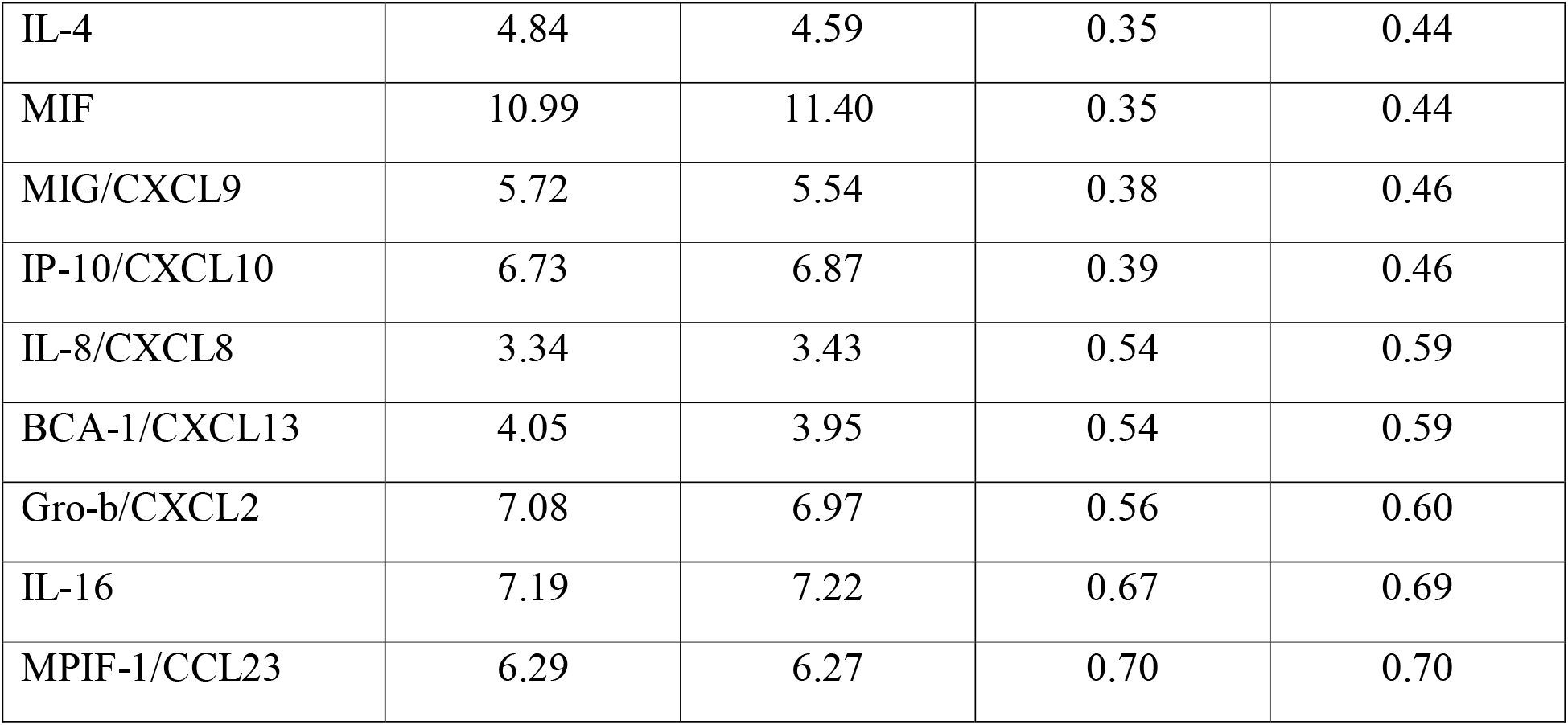
Analysis of 35 cytokines included in the study.

**Supplementa1 Figure 1.**
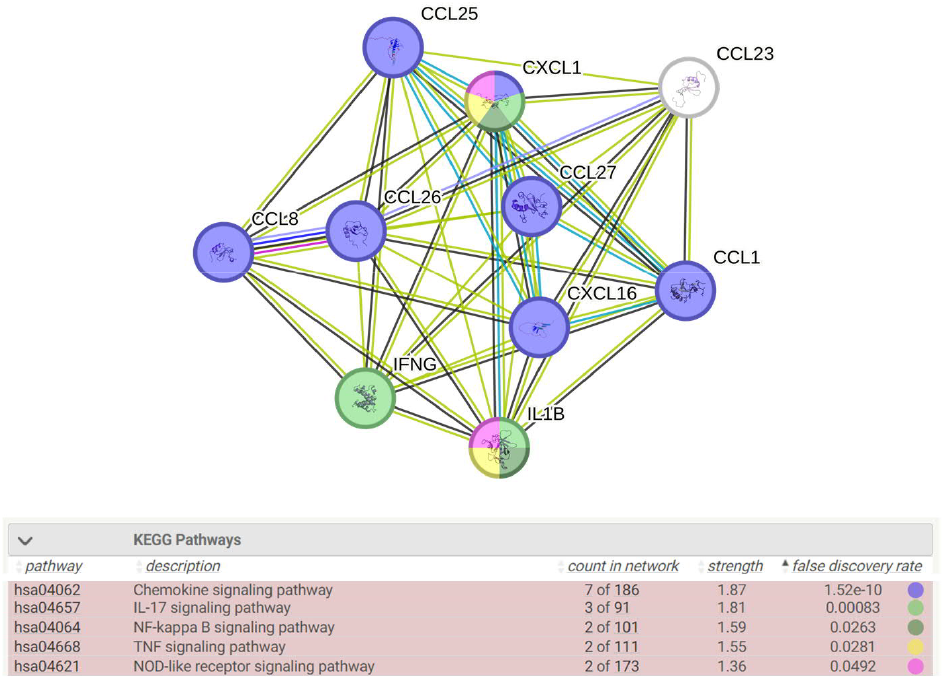
Visual representation of pathways from KEGG database associated with significant cytokines. The signaling pathway analysis was performed using string-db.org. Pathways with strong association by FDR <0.05 are shown. Each network node represents an individual cytokine, and its associated pathways are shown by lines connecting the circles. The colors represent the associated pathways shown in the legend. Images within circles represent cytokine structures.

**Supplemental Figure 2.**
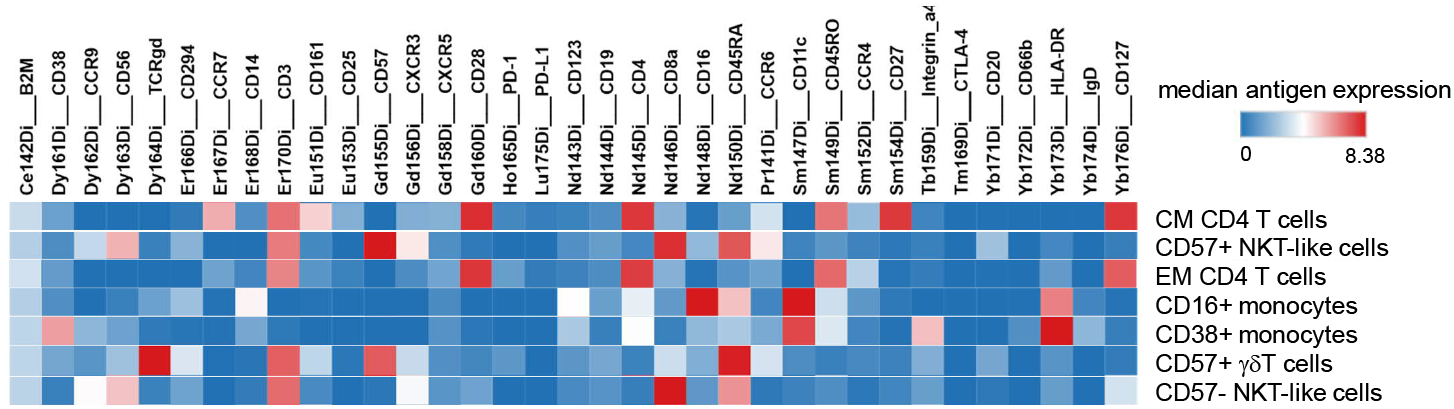
Median intensity of marker expression of clusters shown in **Figure 2B**.

## Author contribution statement

Concept and design: MSv I, DEG

Writing the manuscript: MSvI, JL, DEG

Editing and reviewing the manuscript: All authors

Enrollment of patients: DEG, DH, SC, SB, RK, IP, BS, GAD, YZ, MS

Data abstraction and curation: MSvI, MG, BS

Assay performance: HM-M, FF, PR

Critical analysis and interpretation of data: Ms VI, JAS, JDF, JZ, DEG

Statistical analysis: JL, YX

Administrative support: DEG

## Acknowledgements

The authors thank Ms. Dru Gray for assistance with manuscript preparation.

## Data availability

The datasets used and/ or analyzed during the current study are available from the corresponding author on reasonable request.

